# Changes in postural syntax characterize sensory modulation and natural variation of *C. elegans* locomotion

**DOI:** 10.1101/017707

**Authors:** Roland F. Schwarz, Robyn Branicky, Laura J. Grundy, William R. Schafer, André E.X. Brown

**Affiliations:** European Bioinformatics Institute, Hinxton, UK; MRC Laboratory of Molecular Biology, Cambridge, UK; MRC Clinical Sciences Centre, Imperial College London, London, UK

## Abstract

Locomotion is driven by shape changes coordinated by the nervous system through time; thus, enumerating an animal’s complete repertoire of shape transitions would provide a basis for a comprehensive understanding of locomotor behaviour. Here we introduce a discrete representation of behaviour in the nematode *C. elegans*. At each point in time, the worm’s posture is approximated by its closest matching template from a set of 90 postures and locomotion is represented as sequences of postures. The frequency distribution of postural sequences is heavy-tailed with a core of frequent behaviours and a much larger set of rarely used behaviours. Responses to optogenetic and environmental stimuli can be quantified as changes in postural syntax: worms show different preferences for different sequences of postures drawn from the same set of templates. A discrete representation of behaviour will enable the use of methods developed for other kinds of discrete data in bioinformatics and language processing to be harnessed for the study of behaviour.

**Author Summary:** Technology for recording neural activity is advancing rapidly and whole-brain imaging with single neuron resolution has already been demonstrated for smaller animals. To interpret such complex neural recordings, we need comprehensive characterizations of behaviour, which is the principal output of the brain. Animal tracking can increasingly be performed automatically but an outstanding challenge is finding ways to represent these behavioural data. We have focused on the movement of the nematode worm *C. elegans* to develop a quantitative representation of behaviour as a series of distinct postures. Each posture is analogous to a word in a language and so we can directly count the number of phrases that makes up the *C. elegans* behavioural repertoire. *C. elegans* has a very small nervous system but we find that its behavioural repertoire is still complex. As with human languages, there is a large number of possible phrases, but most are rarely used. When comparing different populations of worms or worms in different environments, we find that the difference between their behaviour is due to a subset of their entire repertoire. In the language analogy, these would correspond to idiomatic phrases that distinguish groups of speakers. A quantitative understanding of the nature of behavioural variation will inform research on the function and evolution of neural circuits.

## Introduction

Animal behaviour can be difficult to study because it involves complex movements that vary continuously in time. This is especially true for limbed animals such as insects and mammals, because they have many degrees of freedom and can adopt an even larger number of different postures. Detailed characterization without explicit posture quantification is possible [1], but researchers commonly use alternative methods for controlling behavioural complexity. In ethology, complexity is often controlled through careful observation to identify a set of behavioural categories that can then be scored to assess differences between individuals, between species, and over time [2]. This has the advantage that observations can be carried out in an animal’s natural habitat where it is able to behave freely. However, the choice of categories may introduce unwanted bias and inter-observer variability can reduce reproducibility [3]. In laboratory-based studies it is possible to impose constraints on the animal, ranging from physically restraining some parts of the body to training an animal to perform a predefined task such as pressing a lever or solving a maze. Constraints reduce measurement dimensionality and can aid interpretation but may also mask variation which may be important for understanding the genetic or neural control of behaviour [4].

Here we adopt an increasingly common [5,6] intermediate approach in which we record animals in a laboratory environment, but with minimal constraints. *Caenorhabditis elegans* is a useful animal for addressing questions of behavioural representation because it has a simple morphology and its movement can be confined to the two-dimensional surface of an agar plate. It has also been characterized exhaustively in several relevant respects: in addition to a complete genome sequence [7], we know the developmental lineage of all of its cells from the single-celled embryo to the adult [8], and it has the most complete connectome—the wiring diagram of its neurons—of any animal [9]. These reference data are powerful in part because of their completeness, making it possible to characterize differences due to genetic, evolutionary, or environmental perturbations systematically. An analogous systematic characterization of *C. elegans* locomotion could also be a useful tool for studying behaviour.

To move on the surface of an agar plate, *C. elegans* propagates bends along its body. It is not clear whether the locomotor wave is generated by a central pattern generator [10], but there is evidence that once generated, waves are driven by proprioceptive coupling between body segments [11]. During navigation, *C. elegans* switches between bouts of forward and backwards locomotion [12] and uses both abrupt and gradual turns during directed behaviours [13,14] and when searching in a relatively featureless environment [15,16]. While performing these manoeuvres, *C. elegans* adopts postures in a low dimensional shape space [17]. However, knowing the dimensionality of the space still leaves open the question of which regions of shape space worms visit and which trajectories they tend to follow. The method we describe here is meant to address this question directly.

We previously described a method of constructing a dictionary of behavioural motifs [18], which are short patterns of repeated motion, and showed that such a dictionary could be useful for identifying and comparing mutant phenotypes in *C. elegans*. However, because motifs can overlap there will be significant redundancy in the representation before completeness is achieved. To solve this problem, we have developed an alternative method of characterising locomotion in terms of a sequence of discrete postures, which serve as building blocks for more complex motifs. We found that 90 distinct postures are sufficient to account for 82% of postural variance. A discrete representation enables efficient enumeration of postural sequences and makes it possible to address questions about behavioural plasticity in response to stimulation and natural variation in spontaneous locomotion. We found that freely moving worms have a core set of frequently used sequences and a much larger set of rarely used sequences. When worms adapt their behaviour in response to environmental conditions such as a lack of food or the presence of an attractive chemical, we find that they adjust the syntax of their locomotion (i.e. their preference for different postural sequences), but that their shapes are still accounted for using the same set of template postures. Similarly, between populations of *C. elegans* isolated from different parts of the world, we find that they have a similar rank-frequency distribution of behaviours, but with significant variations in sequence preference.

## Results

### Ninety template postures capture wild type worm shapes on food

We tracked single worms using a Dino-Lite USB microscope mounted on a motorized stage, as previously described [19]. Worms were segmented from the background using the Otsu threshold and the skeleton was calculated by tracing the midline along the worm body connecting the head and the tail. We then measured the tangent angle at 49 points equally spaced along the worm and subtracted the average angle to achieve a position and orientation independent measure posture [17]. These angles change as the worm bends its body during crawling. Although a worm’s posture varies continuously through time, at any given moment the current posture can be approximated by the closest matching template from a pre-defined set of postures. This is analogous to constructing a histogram, in which discrete bins are used to represent data drawn from a continuous distribution.

The first step was to select a set of template postures that was as small as possible while still having enough elements to capture most of the observed variability. Instead of using uniformly spaced bins to define the templates we used *k*-means clustering, which attempts to minimize the distance between *k* templates (the cluster centres) and the data (the set of input postures measured from wild-type worms crawling on an agar plate seeded with bacterial food). As a distance metric, we used the squared difference between the tangent angle vectors of two worm skeletons. As the number of template postures is increased *(k* in *fc*-means), the R^2^ between the original postures and their nearest neighbours improves (Figure 1A). After 90 postures, the rate of improvement plateaus at a small value (Figure 1A, inset). We therefore used 90 postures for subsequent analyses. The most common postures in this set of templates are sinuous and correspond to different phases of locomotion at different curvatures (Figure 1B). The rare postures include more highly curved shapes that worms adopt during turns as well as straighter shapes that are occasionally observed during dwelling. The full set of 90 postures ordered by frequency is plotted in Figure S1. Using this set of template postures, locomotion can be represented compactly. At any point in time, a worm’s posture is matched to its nearest-neighbour in the set of template postures and so a bout of locomotion can be simply recorded as a series of labels indicating which of the representative postures is closest (Figure 1C).

**Figure 1:**
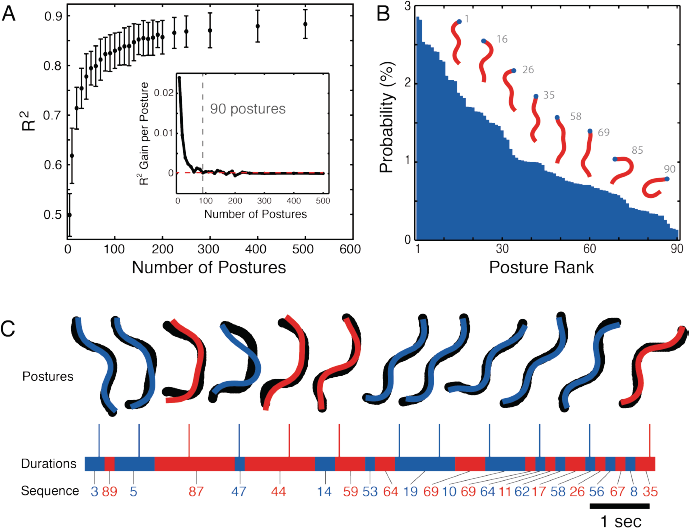
Discrete representation of worm locomotion by clustering. **(A)** As the number of template postures increases, the R^2^ of the fit between original body angles and nearest neighbour matches increases (points show mean ± standard deviation for 50 worms, 15 000 frames each). The inset shows the rate of increase in R^2^ for increasing *k.* The red dashed line is the average increase in R^2^ per posture for *k* = 100 − 500 (2.4 × 10^-4^ per posture). **(B)** Probability of observing each of the 90 template postures. Selected postures are shown as skeletons with the head indicated by the blue dot. The grey label indicates their rank by frequency with 1 being the most common posture. **(C)** Locomotion is represented by a series of discrete template postures. In each frame, the original skeleton (black) is represented by its nearest neighbour (coloured overlay) in the set of 90 template postures. The duration spent in each posture is also recorded and shown as the alternating coloured bands. Each segment of repeated postures is collapsed to a single label and it is this sequence of states that represents locomotion (i.e. {3, 3, 89, 5, 5, 5, 5, 87, 87, 87….} becomes {3, 89, 5, 87}).

### Posture sequences are complex but countable

Repeated patterns of behaviour can now be found simply by finding repeating sequences of postures, including subsequences sampled from longer segments. It is also possible to match similar behaviours performed at different speeds by removing repeated elements from a sequence. For example, if a worm is moving slowly, its nearest-neighbour template posture might not change over several frames leading to a sequence with repeats such as {3, 3, 89, 5, 5, 5, 5, 87, 87, 87, …} (see durations bar in Figure 1C). We reduce this sequence to {3, 89, 5, 87} so that it will be recognized regardless of how slowly the transitions are made. We used this simple kind of non-uniform time warping for all of the sequence results presented in this paper.

In a discrete representation, the behavioural repertoire can be enumerated directly by counting the variety of sequences that are observed. We do this by counting the number of observed *n*-grams. An *n*-gram is any sequence of *n* states, including overlaps. In the sequence {3, 89, 5, 87} the bigrams are {3, 89}, {89, 5}, and {5, 87}, the trigrams are {3, 89, 5} and {89, 5, 87}, and the sequence is itself a single 4-gram. For longer sequences, there are more possible *n*-grams but because of repeats, the number of *unique n*-grams will be less than the total number observed. There is therefore a sublinear growth in the number of unique *n*-grams as more sequences are observed and the degree of this difference is a measure of the structure in the sequence. Figure 2A shows how the number of unique *n*-grams, for *n* = 2-5, grows with total sequence length for the laboratory strain N2. Individual blue lines show data from 100 worms, each recorded for 15 minutes, randomly selected from the total data set of 1262 individuals. Orange lines show accumulation curves for sequences that have been shuffled to destroy temporal order but maintain the relative frequency of postures.

The accumulation curves grow less than linearly, reflecting the fact that worm locomotion is repetitive. It is notable though, that even after over 100 000 postures have been observed, new unique trigrams are still being observed at a rate of approximately 1 in 25 (or about 1 previously unobserved trigram every 12 seconds after a total observation time of 14 hours). This combination of repetitiveness with a large total repertoire can be seen more clearly in a Zipf plot of *n*-gram frequencies (Figure 2B), which shows the frequency of sequences plotted against their rank on a log-log plot. The distribution is heavy-tailed in the sense that there are more high-frequency behaviours than would be expected for an exponential distribution with the same mean (Figure S2). This corresponds to a highly skewed usage of the behavioural repertoire with the top 1% of trigrams being used 32.2 ± 0.6 % of the time during crawling on food (mean ± standard deviation for the blue curves in Figure 2B).

We used the OpenGRM NGram library [20] to model posture sequences as Markov chains, in which the probability of the next posture in the sequence is a function only of the previous *n* – 1 states. In the simplest case of a bigram model, the probability of the next state in the sequence depends only on the current state, but for worm locomotion, a bigram model does not capture the distribution of longer *n*-grams (Figure S3). In contrast, when we fit a trigram model using observed sequences and used it to generate new sequences, the simulated sequences had a similar distribution not only of trigrams, but also 4- and 5-grams (Figure 2A and B, black lines). Increasing *n* to 4 or 5 improves the fit quality, but only modestly (Figure S2). We therefore chose to focus on trigrams for most of the subsequent analysis.

**Figure 2:**
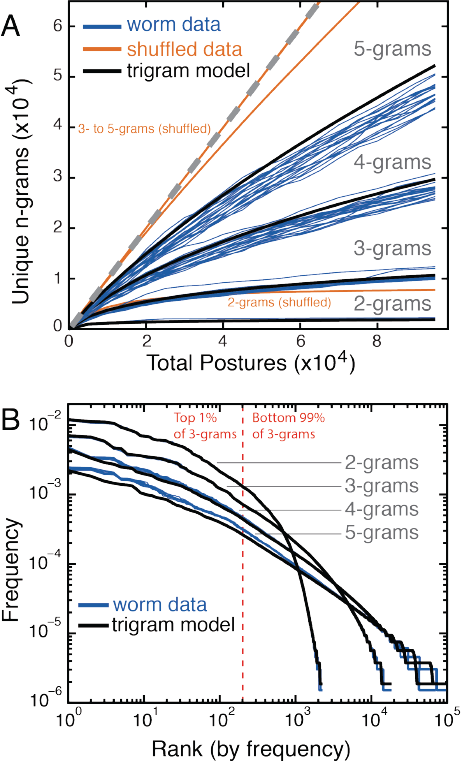
Worm locomotion sequences are complex but can be fit by a trigram model. **(A)** *n*-gram accumulation curves for bigrams (sequences of two postures) to 5-grams (sequences of 5 postures). Over time, as more postures are observed (Total Postures), the number of unique *n*-grams that is observed (analogous to vocabulary size) grows sub-linearly. The grey dashed line has slope 1. Each blue line is calculated from data from 100 randomly chosen worms from the entire data set of 1262 individuals. Orange lines are averages calculated from the same data but with the sequences randomly shuffled. Black lines are averages calculated for data generated from a trigram model of the posture sequences. **(B)** Zipf plot of *n*-grams. The frequency distribution of *n*-grams ranked from most to least frequent. Each blue line is calculated from 400 randomly chosen worms. Black lines are averages calculated for data generated from a trigram model of the posture sequences. The red dashed line indicates the rank that divides the top 1% most frequent trigrams from the remaining 99%.

### Optogenetic activation changes posture *n*-gram frequencies

To determine how worms adapt their behavioural repertoire in response to stimulation, we first used optogenetics to induce behavioural transitions at predictable times in freely crawling worms. We chose two strains for these experiments that produce robust behavioural responses to optogenetic stimulation. ZX46 worms were a useful starting point because they express channelrhodopsin under the control of the *unc-17* promoter in the worm’s cholinergic neurons, and thus activation by blue light results in the simultaneous contraction of body wall muscles and an obvious change in locomotion. Previous experiments on this strain showed that activation leads to a deep dorsal bend [21]. Although dorsal bends are rare under normal circumstances, they are represented in the set of 90 N2 on-food postures and the fit quality during activation actually increases somewhat (Figure 3A). This is because curved postures have a high total variance leading to an R^2^, the fraction of explained variance, near 1. Plots of relative posture probability over time (Figure 3B) show some postures that are overrepresented during the stimulus and a large number of under-represented postures. A rank sum test comparing posture probabilities during and outside of stimulation periods shows that dorsal bends are over-represented (Figure 3C) during stimulation, as previously reported for this strain [21]. The same effect is also seen in the bigram probabilities, although there was insufficient power to detect over-represented trigrams. The under-represented postures are sinuous postures common during normal locomotion. This is due to a decrease in forward locomotion during photoactivation of the cholinergic neurons as revealed by the underrepresented bigrams and trigrams.

**Figure 3:**
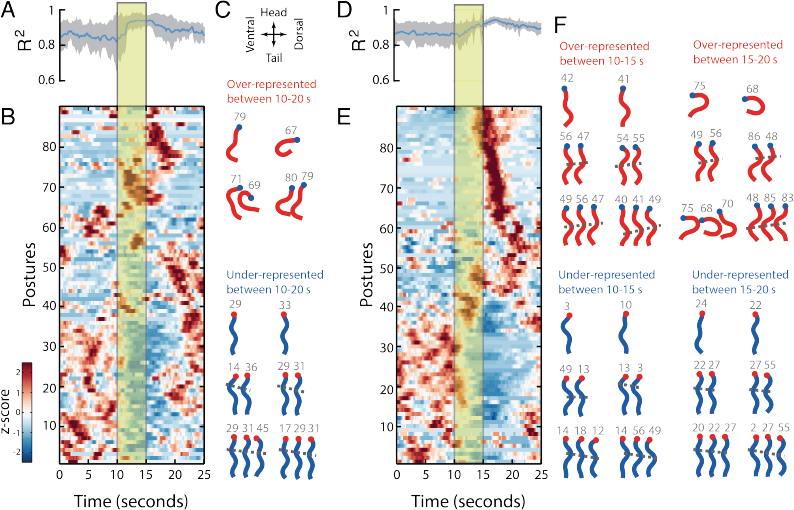
Optogenetic activation of subsets of neurons drives behavioural transitions. The left half of the figure (A-C) shows data from ZX46 worms expressing channelrhodopsin in the cholinergic motor neurons and the right half (D-F) from AQ2026 worms expressing channelrhodopsin in ASH, a neuron that detects aversive stimuli, as well as PVQ and ASI. **(A, D)** R^2^ between original body angles and nearest-neighbour template over time. The blue line is the mean and the gray shading indicates the standard deviation (*n* = 23 in A; *n* = 34 in D). Yellow box indicates light stimulation period. **(B, E)** The relative probability of observing each posture over time. Colour indicates whether a posture is over-represented (red) or underrepresented (blue) at each time. Postures with similar temporal profiles are clustered independently in B and E for visualization. **(C, F)** The top 2 *n*-grams that are most significantly overrepresented (red) or underrepresented (blue) during the indicated time periods for *n* = 1-3 for each strain. Grey numbers show the row number for each posture in the corresponding heatmap. Dots indicate worm head, dorsal and ventral sides are as indicated by the arrows. The gray dashed lines approximately connect body bends from one posture to the next and can be used to distinguish forward from reverse locomotion. Lines with a negative slope indicate body bends propagating backwards and therefore forward locomotion. Lines with a positive slope show reversals.

Photoactivation of AQ2026 worms, which express channelrhodopsin under the control of the *sra-6* promoter, reliably induces an escape response through the activation of the ASH sensory neurons [22,23]. The behaviour is readily observed, but is more naturalistic than the deep dorsal bends seen in ZX46 worms since it occurs through activation of a sensory neuron that normally detects aversive stimuli and elicits an escape response. Again, the discrete representation fits the induced postures well (Figure 3D). The plot of posture probability over time shows both over- and under-represented postures during and after stimulation (Figure 3E). Escape responses in *C. elegans* typically consist of a reversal followed by a turn, which we can detect by splitting the analysis into three parts: outside stimulus, during stimulus, and 0-5 seconds after stimulus. During stimulation the bigrams and trigrams representing reversals are significantly more likely while forward locomotion is less likely (Figure 3F) and in the 5 seconds after stimulus, ventral turns and reversals are more likely while forward locomotion is still supressed.

### Worms change their postural syntax in different environments

We next recorded worm locomotion in different environments to see how they adapt their behavioural repertoire in response to variation in sensory stimulation. To add to the data for worms on bacterial food, we also recorded N2 worms crawling on an agar surface with no food, and with no food but in the presence of the chemo attractant benzaldehyde [24]. N2 animals rarely leave a patch of food [25] and spend some time dwelling leading to a confined path (Figure 4A). In the absence of food, worms show a clear switch in behaviour, spend more time roaming and initiate a local search behaviour consisting of reversals and turns. After longer periods of starvation, they show increasingly directed locomotion [15,16]. During chemotaxis off food, their change in behaviour compared to worms off food with no chemo attractant is more subtle; they still spend most of their time moving, but their paths are less random as they will, on average, move up the gradient of attractant [13,14].

**Figure 4:**
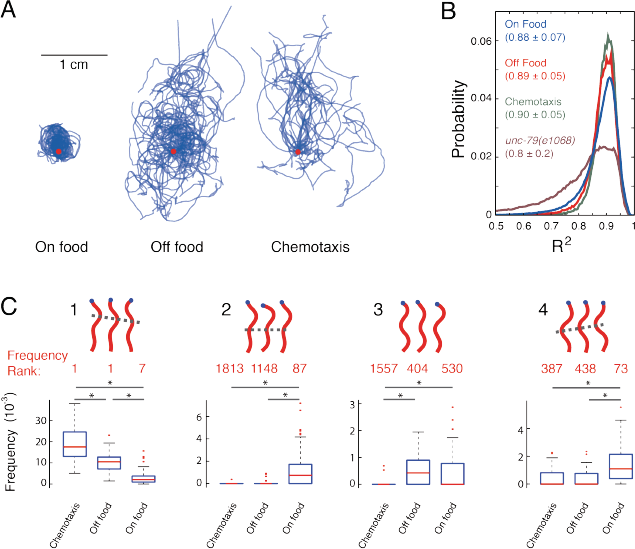
Worm behaviour is compositional across different conditions. **(A)** Worm tracks in different conditions: on food (n = 34), off food (n = 23), and off food with an attractant (benzaldehyde) (n = 25). Tracks for different worms start at the red dot and are aligned with their long-axis arranged vertically. Worms on food explore the region of the food but rarely leave, while off food they explore a larger area. In the presence of an attractant, worms explore a large area, but in a more directed way. **(B)** A plot shows the distribution of R^2^ values for worms in the different conditions. The fit quality is no worse for worms off food or performing chemotaxis even though the postures are derived only for worms on food. For comparison, the distribution of fits is also shown for an uncoordinated mutant *unc-79(e1068)* that is poorly fit using the wild type on food postures. Numbers in the legend are mean ± standard deviation of the R^2^ distributions. **(C)** Comparisons between the frequencies of four selected trigrams in the three conditions. The rank of each sequence in the repertoire of worms in each condition is shown in red. * indicates *p* < 4.1 × 10^-4^, chosen to control the false discovery rate at 5% across multiple comparisons.

We used the same set of 90 templates derived for N2 animals on food to fit the shapes of worms off food and during chemotaxis. The fit quality is no worse in either of the off food conditions, indicating that there is not an increase in outlying postures that are not captured by the template set (Figure 4B). In fact, the poorly fit tail of the distribution is reduced off food compared to on food. The better fit may be due too noisier skeletonization of worms imaged on food and partly because worms off food are less likely to adopt some postures characteristic of pauses that are among the worst fitting postures (Figure S4). As a positive control, we also measured the distribution of fit qualities for a visibly uncoordinated mutant *unc-79(e1068)* and found a clear increase in outlying postures not captured by the wild type on food template postures.

We also tested for differences in *n*-gram frequencies that worms performed in the different conditions by pooling their sequences and getting the complete set of unique *n*-grams observed across the three conditions, for *n* = 1-3. We counted the number of occurrences of each *n*-gram in each condition and used a rank sum test to find sequences that were used more or less frequently in one of the conditions. We excluded *n*-grams that were observed less than 5 times in total in the two conditions being compared. Note that we did not exclude behaviours that were rare in only one of the conditions. For example, an *n*-gram with a count of 10 in one condition and 0 in another would still be included in the comparison. We corrected for multiple comparisons using an adjusted p-value threshold calculated to control the false discovery rate at 5% [26]. Using this procedure, we found that for the on food vs. off food comparison 377 trigrams were significantly more common in one condition, whereas for the subtler off food vs. chemotaxis comparison we found 11. These differences in trigram frequencies are not entirely explained by differences in posture preference, since in shuffled sequences that preserve posture frequency differences, only 2 trigrams were found to be significantly different between the on and off food conditions and none were significantly different between the off food and chemotaxis conditions. Four examples of distinguishing trigrams are shown in Figure 4C. The first one is a bout of forward locomotion that is common in all conditions, but progressively less common in worms performing chemotaxis, crawling off food, and crawling on food. This is consistent with expectations for the persistence of locomotion in these conditions. The second trigram starts with the same posture as the first, but it appears in a different context. Instead of appearing during forward locomotion, the posture appears during a pause with a head swing. This change in context makes trigram 2 much less common in the off food conditions even though trigrams 1 and 2 share a posture in common. This implies that behavioural adaptation is at least partly characterized by shifts in postural syntax. Trigram 3 shows that behaviours that are rare in all conditions can still show significant differences in frequency and trigram 4 shows an example of a reversal which is more common on food than in either of the off food conditions, consistent with worms showing more persistent forward locomotion when searching off food.

### Natural variation in behaviour is captured by a subset of posture *n*-grams

In addition to environmental variation, we also quantified the behavioural variation that arises over short evolutionary times between different populations of the same species. We tracked 17 strains of *C. elegans* isolated from different regions of the world and quantified their behaviour using the discrete representation. We found that the template postures derived from N2 worms on food could also fit the wild isolate data and that the heavy-tailed frequency distribution of trigrams is observed in all strains (Figure 5A). The distribution is skewed with the top 1% of trigrams being used 38 ± 4 % of the time during crawling (mean ± standard deviation, *n* = 17 strains).

**Figure 5:**
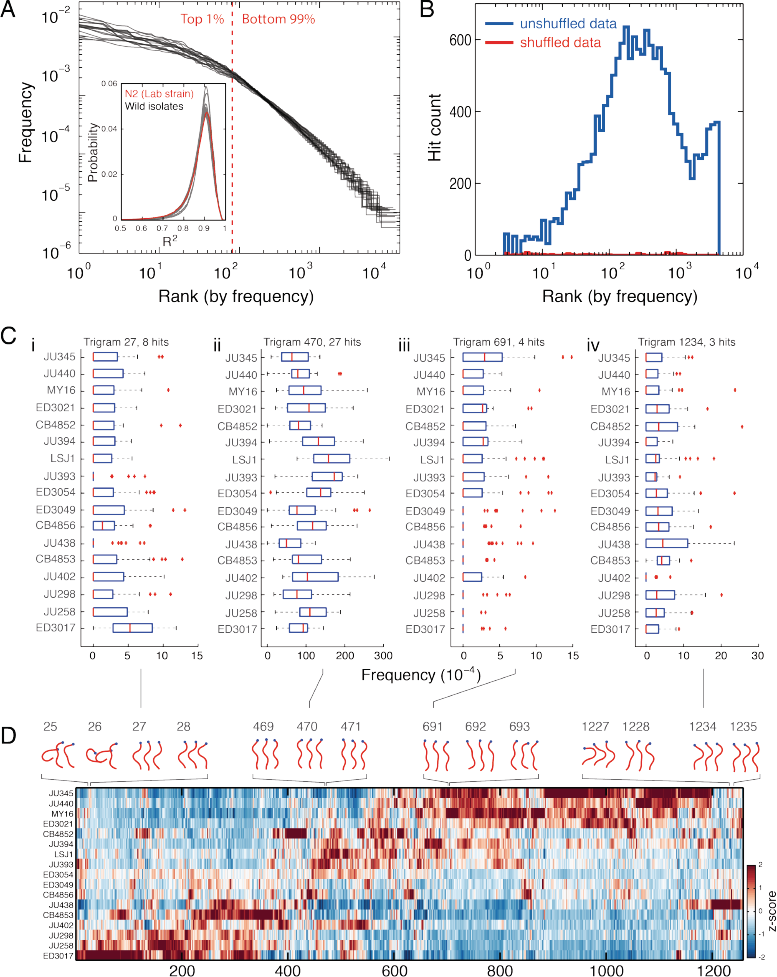
Form of trigram frequency distribution is conserved across wild isolates despite differences in preference for particular sequences. **(A)** Zipf plots for 17 wild isolates of *C. elegans* all show a heavy tailed distribution similar to N2 worms. The red dashed line shows the rank dividing the top 1% of trigrams in the repertoire from the remaining 99%. The inset shows the distribution of R^2^ values for fits between 90 N2-derived templates and the original postures for the 17 wild isolates as well as for N2 itself for comparison. The overall mean of the distributions is 0.86 with standard deviation 0.08. **(B)** Histogram of hit counts (not hit density) versus frequency on a log scale. Hits are trigrams with significantly different frequencies in at least one pairwise rank sum test comparing strains. Note that the bin widths increase to be constant on a log-scale. Blue line is the original data and the red line is for trigrams derived from the same data after random shuffling of the sequences. **(C)** Box plots of trigram frequencies for 4 trigrams showing hits in several strains. The trigram number indicates its column in the heatmap in D. The p-value threshold was set at 1.1 × 10^-4^ to control the false discovery rate at 5% across comparisons. **(D)** A heatmap showing the relative frequency of all of the trigrams with significantly different frequencies in at least one strain comparison. Strains and trigrams have been hierarchically clustered. Sample trigrams are plotted above the heatmap with their columns indicated by the gray numbers.

We performed rank sum tests between all strain pairs to find trigrams with significant frequency differences using a *p*-value threshold adjusted to control the false discovery rate at 5% across all comparisons. There are trigrams with significant differences across the frequency spectrum; in fact most behaviours with a significant frequency difference are rare in at least one of the strains where the behaviours were found as a hit (Figure 5B). Not all of the hits counted in the histogram in Figure 5B correspond to unique trigrams because some trigrams had significantly different frequencies in more than one strain (Figure 5C). The number of *unique* trigrams that are different in at least one strain is 1258, which is approximately 5% of the 2.3 × 10^4^ distinct trigrams observed in the full wild isolate data set. The relative frequencies for these 1258 trigrams are shown in a heatmap in Figure 5D. Trigrams 25 and 26 show dorsal turns. 27 and 28 are also bent more dorsally than ventrally, but 27 represents forward locomotion while 28 represents a reversal. 469-471 are all bouts of forward locomotion, 691-693 are pauses with relatively straight postures. 1227 is a ventral turn and 1234 and 1235 are reversals.

## Discussion

We have developed a simple representation of behaviour as a series of discrete postures to systematically enumerate the behavioural repertoire of *C. elegans* during locomotion. The concept of a complete description of behaviour was motivated in part by similar efforts devoted to other aspects of worm biology, most notably the genome, cell lineage, and connectome. However, the repertoire of behavioural sequences cannot be said to be complete in a similar way. Although we have a complete record of the observed sequences, accumulation curves suggest that more recordings would continue to reveal novel sequences, even when considering behaviours with durations of about a second in an animal with a simple morphology in a relatively uniform environment. A notable feature of the observed repertoire is the simultaneous presence of repetition (the heavily-used top 1%) and variety (the large set of relatively rarely used sequences). This feature may result from two demands on an animal’s behavioural repertoire: locomotion should be efficient in a given environment but must also be flexible to deal with unexpected conditions or to respond to new sensory information. The repertoires of 17 wild isolates of *C. elegans* have a similar rank-frequency distribution and we hypothesize that such heavy-tailed distributions will generalize beyond nematodes and will in fact be a widespread characteristic of animal behavioural repertoires.

Some of the rarely observed sequences in the repertoire are surely due to measurement noise, but since the frequency of some rare behaviours is modulated in response to sensory stimuli (for example behaviour 3 in Figure 4C), measurement noise is not the only factor. It should also be emphasized that although the repertoire is large in this representation, that does not preclude the existence of an alternative representation that will reveal an underlying simplicity. Indeed, seemingly simple nonlinear systems are well known to be able to produce complex dynamics [27] and the search for such models in the context of animal behaviour remains an exciting challenge [28-30]. The observation that only 5% of the trigram repertoire is modulated by natural variation between wild isolates (Figure 5D) may be a reflection of a small number of behavioural modules that can be adjusted genetically within the constraints of natural selection [31,32].

New imaging technology has made it possible to record neural activity from the majority of *C. elegans* neurons simultaneously in restrained animals [33] and during locomotion [34]. The usefulness of this data will depend to some extent on how the corresponding behavioural data is represented and quantified. This is also the case for optogenetically induced behavioural changes as reported here and many times previously in *C. elegans* (e.g. [35–39]). Recent work in *Drosophila* larvae suggests that an unsupervised detection of locomotion patterns can be informative [40]. The discrete approach that we report here is also unsupervised in the sense that repeated patterns of behaviour are detected based on locomotion patterns observed within strains without initially using information about different classes (environmental conditions or strains). A conceptually similar approach has proven to be useful for quantifying drug-induced behaviour in mice [41]. It is likely that no behavioural representation will be optimal for all studies, but that different representations will provide complementary insight. The advantages of trigram counting are that it is relatively unbiased, can reliably connect behaviours performed at different speeds, provides a complete record of all of the observed behaviours (up to the resolution of the template postures), and that it can be summarized intuitively as frequencies of short behaviours that can be directly visualized. For some applications the variance that is missed by the discrete templates may be critical. One option would be to increase the number of templates, but it may also be that a different representation would be called for. Another limitation of the discrete representation as presented here is that we have not used the information contained in the duration that worms spend in each posture. The time that worms spend performing each behavioural sequence is relatively straightforward to incorporate and would increase the power of the representation to resolve differences in dynamics between strains.

In the wild, animals must adapt to a variety of environments that can change unpredictably. One strategy for dealing with this complexity is to use a finite set of behavioural primitives that can be composed in different sequences to generate adaptive responses [42]. This is precisely the kind of variation that we observed for worms changing their behaviour in response to new environments (Figure 4). Using the same set of postures, they change their behaviour at the level of postural syntax. Compositionality in behaviour also suggests a mechanism for generating complex behaviours by combining primitive actions. Hierarchical composition has long been recognized as a principle of animal behaviour [43,44] and has also been proposed as an efficient representation for learning in robotics [42]. The possible connection between motor learning and compositionality raises the interesting question of how the repertoire observed in adults is acquired and modified during development.

## Materials and Methods

### Worm Tracking

Locomotion data was recorded for single worms as previously described [19]. For on food tracking, single worms from each strain were picked to the centre of a 25 mm agar plate with a spot of *E. coli* OP50 at the centre, allowed to habituate to the new plate for 30 minutes, and then recorded for 15 minutes. The worm side (whether it was on its left or right side) was manually annotated using a stereomicroscope before transferring plates to the tracking microscope. In addition to the previously recorded N2 and DR96 *unc-76(e911)* data, we used the following wild isolate strains: CB4852, CB4853, CB4856, ED3017, ED3021, ED3049, ED3054, JU258, JU298, JU345, JU393, JU394, JU402, JU438, JU440, LSJ1, and MY16. For experiments off food, worms were briefly transferred to plates without food and then picked to the centre of a 55 mm agar plate and recorded immediately. Recording was stopped once worms reached the edge of the plate. To maximize the time of recording, worms in the chemotaxis assays were picked to a location near the edge of a 55 mm agar plate opposite the location of the attractant spot. 1 μl of the attractant, benzaldehyde (diluted 1:100 in EtOH), was added to the plate 10 mm from the edge.

### Optogenetics

The strains used for optogenetics were ZX46 *zxIs6 [punc-17::ChR2(H134R)::yfp]* V [21] and AQ2026 *ljIs105 [sra-6::ChR2::yfp, unc-122::gfp]*[22]. Worms were transferred to retinal plates at the L4 stage, left at 20°C, and assayed either 24 hours later as adults (AQ2026) or in the next generation (ZX46). Retinal plates were made by seeding NGM plates with OP50 mixed with all-trans retinal (100mM in ethanol) in a ratio of 1000:4. For channel rhodopsin activation, worms were illuminated for 5 seconds with 1 mW/mm^2^ blue light (440-460 nm) using LEDs (Luxeon III LXHL-PR09 from Lumileds, CA) controlled by a Mindstorms LEGO Intelligent NXT Brick. Each worm was stimulated three times, with an interstimulus interval of 30 seconds.

### Posture Discretization

The skeleton of worms in each frame was found after segmentation by tracing the midline between the two points of highest curvature (the head and tail) of the largest connected component in the image (the worm body) and down sampling to 49 equally spaced points [19]. The skeletons were then converted to a position and orientation independent representation by measuring the angle between each of the 49 skeleton points and subtracting the mean angle [17]. Missing skeleton angles were linearly interpolated. Missing data resulted from camera dropped frames, stage motion, or failure to skeletonize because of poor segmentation or coiled worms. On average 13 ± 9% (mean ± standard deviation) of frames cannot be segmented with each of the three causes contributing approximately equally to missing data. 5000 skeletons were randomly selected from each of 20 randomly selected N2 worms recorded crawling on food. Worms were flipped so that all worms appear to be on their right side when viewed on the page. This means that dorsal and ventral turns are consistent across individuals during analysis. This set of skeletons was then clustered using *k*-means clustering to identify a set of representative template postures. *k* was varied from 5 to 500 to determine the value for *k* after which increasing *k* only marginally improves the quality of the fit. *k* was chosen as 90 for all subsequent analysis.

For each worm video, we down sampled the data from 30 frames per second to 6 frames per second and then found the best matching template in each frame of the down sampled data. The best match was determined in each frame by finding the template posture with the minimum squared difference to the skeleton in that frame. Locomotion was then represented as a series of state labels between 1 and 90. Repeated states were collapsed to a single state (e.g. {1, 1, 2, 2, 2, 5} would become {1, 2, 5}).

### Posture Probability Analysis

For the optogenetics data, the frequency of each posture over time is averaged over three trials per worm and further averaged over the individuals in each experiment (*n* = 26 for ZX46 and *n* = 34 for AQ2026). Each row was smoothed with a moving average filter with a width of 1.3 seconds. The smoothed frequencies of each posture over time were then normalized by subtracting the mean and dividing by the standard deviation of each row to show relative up- and down-regulation over time. To aid visualization, the rows were then clustered vertically so that rows with similar temporal profiles were near each other. The clustering was agglomerative hierarchical clustering using Euclidean distance and average linkage. The clustering was performed independently for the data in Fig. 3B and E so the rows in each plot do not correspond to the same postures.

### n-gram model

The *n*-gram language model analyses were implemented in C and Python using the OpenFST [45] and OpenGRM *n*-gram [20] libraries. Postural sequences from N2 worms on food were compiled into finite-state archives (FAR) and standard *n*-gram count models were built for *n* = 1-5 and normalized into probabilistic models without additional smoothing (see e.g. [46]). For the simulation studies and for each n, 1000 random sequences were generated from the normalized finite-state models. Beginning at the start state an outgoing arc is chosen at random proportionally to the log-probability associated with the arc, the postural state is emitted and the process continues until an end state transition is reached.

### n-gram Statistics

The total set of unique *n*-grams was determined for each strain or condition (for the on food, off food, and chemotaxis set, the total set contained 14 209 trigrams and for the 17 wild isolates it contained 22 638 trigrams). Counts of unique *n*-grams in each individual were then used to compare strains or conditions to each other. For each comparison, any *n*-gram that was observed a total of 5 times or less in both groups being compared was ignored. For the remaining *n*-grams, counts were compared using a rank sum test and *p*-value thresholds were chosen using the Benjamini-Yekutieli procedure to control the false discovery rate at 5% [26].

## Acknowledgements

We thank Mario de Bono and Kate Weber for providing us with the wild isolates used in this study and for and useful discussions and Alexander Gottschalk for providing ZX46 worms. This work was funded by the Medical Research Council through grants MC-A022-5PB91 to WRS and MC-A658-5TY30 to AEXB and supported in part by the National Science Foundation under Grant No. PHYS-1066293 and the hospitality of the Aspen Center for Physics.

---

**Figure S1:**
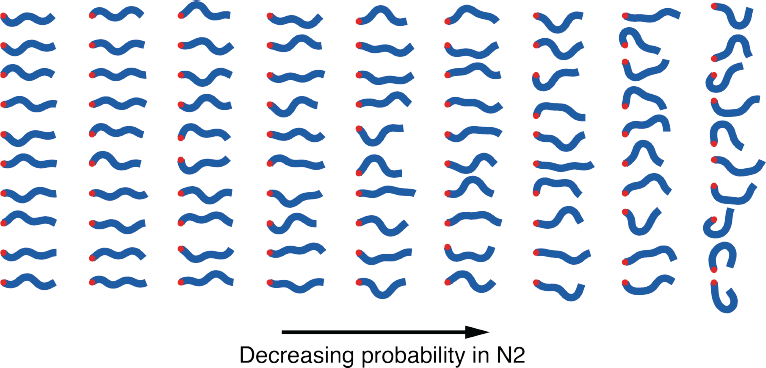
The complete set of 90 posture templates derived for N2 animals crawling on food using *k*-means clustering. The postures are ordered from most frequent (top left) counting down each column in turn to the least frequent (bottom right).

**Figure S2:**
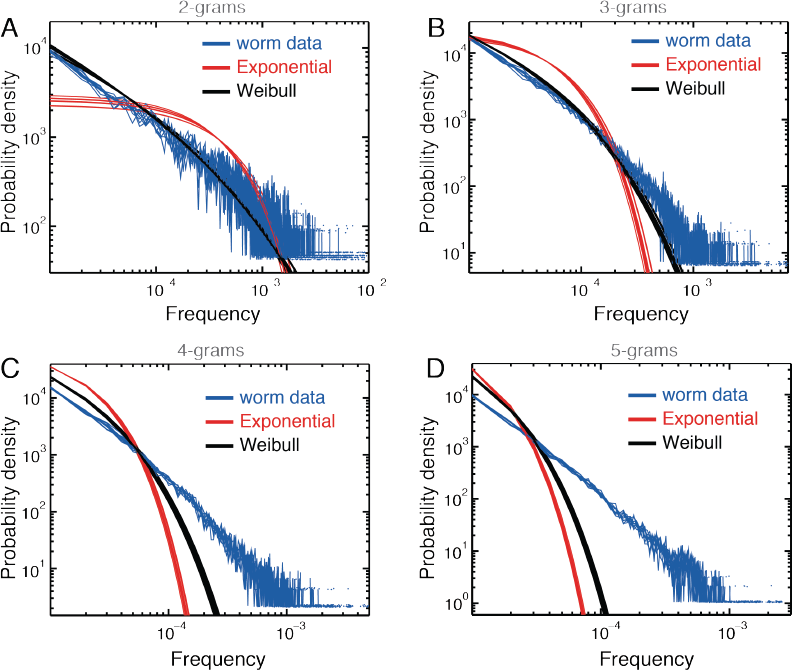
Frequency distribution of *n*-grams compared to exponential distributions and Weibull distributions with parameters set by maximum likelihood estimation. Blue curves are each calculated for 400 worms resampled with replacement from the total N2 data set. Each of the 10 curves is fit separately to an exponential (red curves) and a Weibull (black curves) distribution. A-B show results for *n* = 2 – 5.

**Figure S3:**
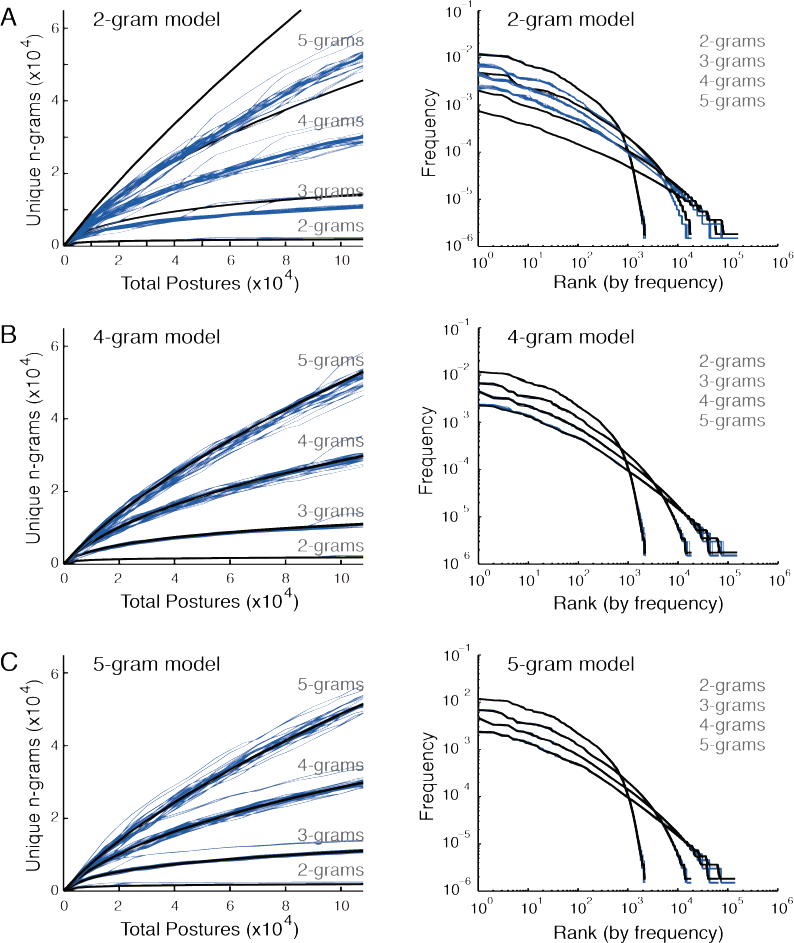
Accumulation curves (left column) and Zipf rank-frequency plots (right column). Blue lines show data from different randomly selected sets of N2 worms crawling on food (100 worms selected for each curve). Black lines show the corresponding results for data generated from *n*-gram models with different values of *n* shown in A-C.

**Figure S4:**
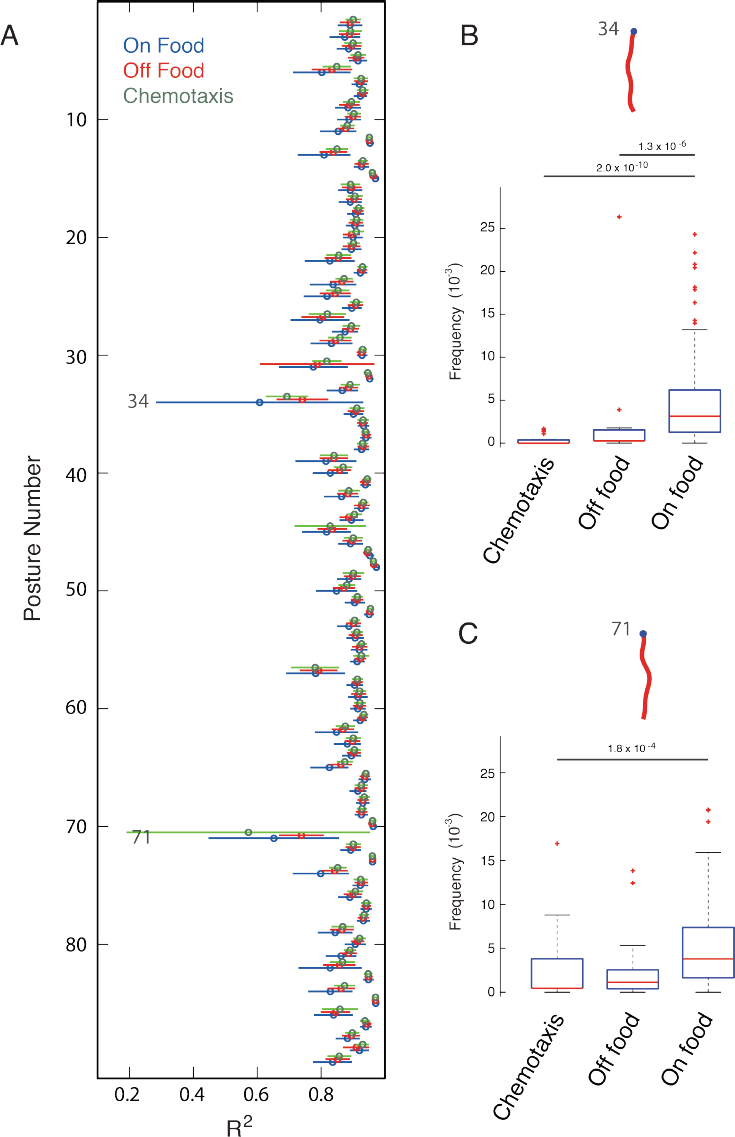
(A) R^2^ plotted for each posture individually (mean ± standard deviation) for data collected on food, off food, and during chemotaxis. The mean fit quality is consistently slightly lower on food than in the two off food conditions, consistent with slightly noisier segmentation when worms are on a patch of food. Furthermore, the two postures with the lowest R^2^ values (34 and 71) are used less frequently in the off food conditions than when worms are on food **(B)**.

